# Altered directed functional connectivity of the right amygdala in depression: high-density EEG study

**DOI:** 10.1101/620252

**Authors:** Alena Damborská, Eliška Honzírková, Richard Barteček, Jana Hořínková, Sylvie Fedorová, Šimon Ondruš, Christoph M. Michel, Maria Rubega

## Abstract

The cortico-striatal-pallidal-thalamic and limbic circuits are suggested to play a crucial role in the pathophysiology of depression. Stimulation of deep brain targets might improve symptoms in treatment-resistant depression. However, a better understanding of connectivity properties of deep brain structures potentially implicated in deep brain stimulation (DBS) treatment is needed. Using high-density EEG, we explored the directed functional connectivity at rest in 25 healthy subjects and 26 patients with moderate to severe depression within the bipolar affective disorder, depressive episode, and recurrent depressive disorder. We computed the Partial Directed Coherence on the source EEG signals focusing on the amygdala, anterior cingulate, putamen, pallidum, caudate, and thalamus. The global efficiency for the whole brain and the local efficiency, clustering coefficient, outflow, and strength for the selected structures were calculated. In the right amygdala, all the network metrics were significantly higher (p<0.001) in patients than in controls. The global efficiency was significantly higher (p<0.05) in patients than in controls, showed no correlation with status of depression, but decreased with increasing medication intake (R^2^ = 0.59 and p = 1.52e − 05). The amygdala seems to play an important role in neurobiology of depression. Practical treatment studies would be necessary to assess the amygdala as a potential future DBS target for treating depression.

---

Affective disorders belong to the most common and most serious psychiatric disorders ^1^. A crucial role of the cortico-striatal-pallidal-thalamic and limbic circuits in the neurobiology of depression was repeatedly reported ^2 3 4^. Magnetic resonance imaging, functional magnetic resonance imaging (fMRI), magnetoencephalographic, and electroencephalographic (EEG) studies have confirmed that depressive patients show structural impairments and functional disbalances of brain networks that involve structures engaged in a) emotions, i.e. amygdala, subgenual anterior cingulate, caudate, putamen and pallidum ^5 3 6 7 8 9 10 11 12^; b) self-referential processes, i.e. medial prefrontal cortex, precuneus, and posterior cingulate cortex ^13 14^; c) memory, i.e. hippocampus, parahippocampal cortex ^15^; d) visual processing, i.e. fusiform gyrus, lingual gyrus, and lateral temporal cortex ^16^; and e) attention, i.e. dorsolateral prefrontal cortex, anterior cingulate cortex (ACC), thalamus, and insula ^17 10 11 12^. Moreover, post-mortem morphometric measurements revealed smaller volumes of the hypothalamus, pallidum, putamen and thalamus in patients with affective disorders ^18^.

Many depressive patients fail to respond to pharmacological therapy resulting in 1 – 3% prevalence of treatment-resistant depression (TRD) ^19^. One of the newest therapeutic approaches for TRD is an invasive direct electrical stimulation of relevant deep brain structures ^20^. Both unipolar and bipolar depression patients might benefit from deep brain stimulation (DBS) treatment ^21^, although an optimal approach, including selection of an optimal target structure, has yet to be established. Selection of the brain structures, that are currently being tested as DBS targets for treating depression ^20^, is mostly supported with the evidence from lesional ^22 23^, animal ^24 25 26 27 28 29 30^, and neuroimaging ^31 32 33 34 35 36 37 38^ studies. The latter approach provides evidence from a network perspective ^39 40^ showing dysbalances in the intrinsic functional architecture of the brain. During a resting state, patients with depression as compared to healthy controls show hyperconnectivity within the default mode network ^13 33 38^, hypoconnectivity within the frontoparietal network ^41 42^, hyperconnectivity between the default mode and frontoparietal networks ^43^, and dysbalances in connectivity within the salience ^44 45^ and dorsal attention ^46^ networks. Functional connectivity anomalies between the hippocampus, cortical and subcortical regions ^47^ similar to those observed in humans with depression, were also observed in a genetic rat model of major depression. The pathophysiological basis of depression, however, still remains incompletely understood. Particularly, better understanding of the connectivity properties of deep brain structures potentially implicated in DBS treatment could have an important value.

Neuroimaging techniques, such as fMRI and EEG, allow to investigate the integration of functionally specialized brain regions in a network. Inferring the dynamical interactions among simultaneously recorded brain signals can reveal useful information in abnormal connectivity patterns due to pathologies.

The connectivity studies based on fMRI are usually based on correlation analyses without providing knowledge about the direction of the information flow between the examined regions. Understanding the directionality is, however, crucial when searching for suitable DBS targets for treating TRD, because the antidepressant effect of DBS treatment might be caused by changes in the activity of remote structures that receive inputs from the stimulated region. For example, it has been hypothesized that DBS applied in the nucleus accumbens might influence the activity in the ventral (subgenual ACC, orbitofrontal and insular cortices) and dorsal (dorsal ACC, prefrontal and premotor cortices) subnetworks of the depression neurocircuitry ^48^. Causal link between a functional inhibition of the lateral habenula and reduction of the default mode network hyperconnectivity was shown on a rat model of depression ^30^, which might explain the therapeutic effect of the lateral habenula DBS in TRD patients ^49^. In other words, the functional inhibition of a deep brain structure via DBS might cure depression through reduction of the hyperconnectivity in the large-scale brain network. Another example of a particular role of the stimulated structure in the large-scale neural communication is the ACC, whose possible integrative role in cognitive processing ^50 51^ might explain the most recently reported high efficacy of DBS to subgenual ACC in treating depression ^52^.

The growing interest in investigating the dynamical causal interactions that characterize resting-state or task-related brain networks has increased the use of adaptive estimation algorithms during recent years. Particularly, Granger causality based on adaptive filtering algorithms is a well suited procedure to study dynamical networks consisting of highly non-stationary neural signals such as EEG signals ^53 54^. The adaptive filtering enables to deal with time-varying multivariate time-series and test direct causal links among brain regions. A signal *x* is said to Granger-cause another signal *y* if the history of *x* contains information that helps to predict *y* above and beyond the information contained in the history of *y* alone ^55^.

Aberrant functional EEG-based connectivity in depressive patients was reported in studies where network metrics were computed directly between sensor recordings ^56 57 58 59 60 61^. Since each EEG channel is a linear mixture of simultaneously active neural and other electrophysiological sources, whose activities are volume conducted to the scalp electrodes, the utility of such observations on the sensor level is limited ^62 63^. This limitation is particularly remarkable in connectivity studies which aim to identify the real active relations between brain regions. Connectivity analysis performed in the source space enables to partially overcome this issue ^62^. Indeed, Partial Directed Coherence estimators do not take into account zero-lag interactions that describe the instantaneous propagation of activity, considering the zero-phase connectivity as noise added to lagged connectivity patterns of interest. For this reason, directed functional connectivity analysis based on electrical source imaging proved to be a promising tool to study the dynamics of spontaneous brain activity in healthy subjects and in various brain disorders ^64 65 66^. Despite this fact, the electromagnetic imaging has not been yet used in patients with depression to study the directed connectivity of resting-state networks.

In the current study, we explored the directed functional connectivity at rest in depression using high-density EEG. We computed the Partial Directed Coherence on the source EEG signals focusing on the role of the amygdala, anterior cingulate, putamen, pallidum, caudate, and thalamus in large-scale brain network activities. We hypothesized that the resting-state directed functional connectivity in these deep brain structures might be disrupted in patients with depression compared to healthy controls.

## Results

In line with the aim of the study we focused on resting-state electrophysiological activity of twelve regions of interest (ROIs) of selected deep brain structures. Further details on results on the ROIs of the whole brain are reported in the Supplementary Information.

### Power spectra

We found an overall increase in power in theta and alpha frequency bands in patients compared to controls at both the *population* and *single-subject* levels. At the *population* level, significantly higher power (p<0.05) in patients was found in all investigated subcortical regions in both frequency bands (see Figure 1). At the *single-subject* level, a significantly higher power (p<0.05) in patients than in controls was observed in the [4-12] Hz frequency range bilaterally in the thalamus, pallidum, putamen, and caudate. Moreover, a significant left-lateralized power increase (p<0.05) in patients vs controls was observed in the anterior cingulate and amygdala in this frequency range (see Figure 2b).

**Figure 1.**
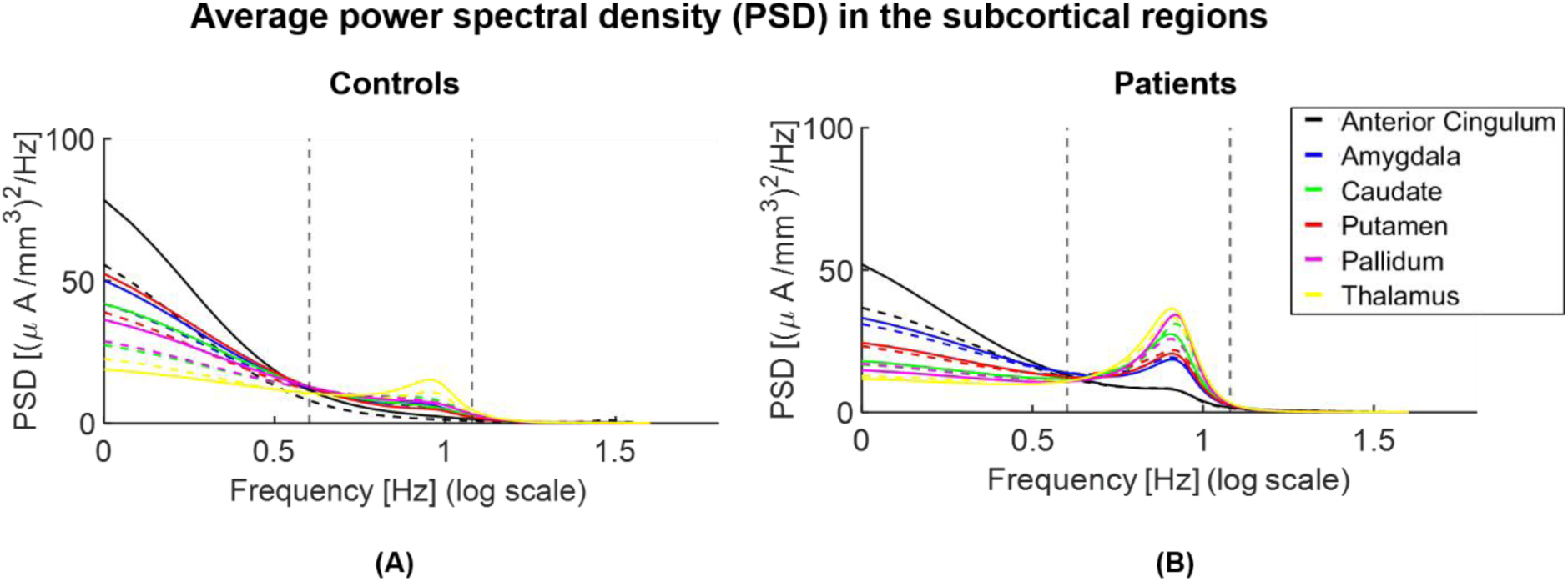
Parametric power spectral density (PSD) of the *population subjects* representing controls (A) vs patients (B) in the subcortical regions of interest. Power significantly increases within the interval [4-12] Hz (indicated with vertical dashed lines) in theta ([4-8] Hz) and alpha ([8-12] Hz) bands and decreases in delta ([1-4] Hz) and beta ([12-18] Hz) bands in patients compared to controls (p<0.05) in the subcortical regions of interest. Continuous and dashed lines indicate the results for structures in the right and left hemispheres, respectively.

**Figure 2.**
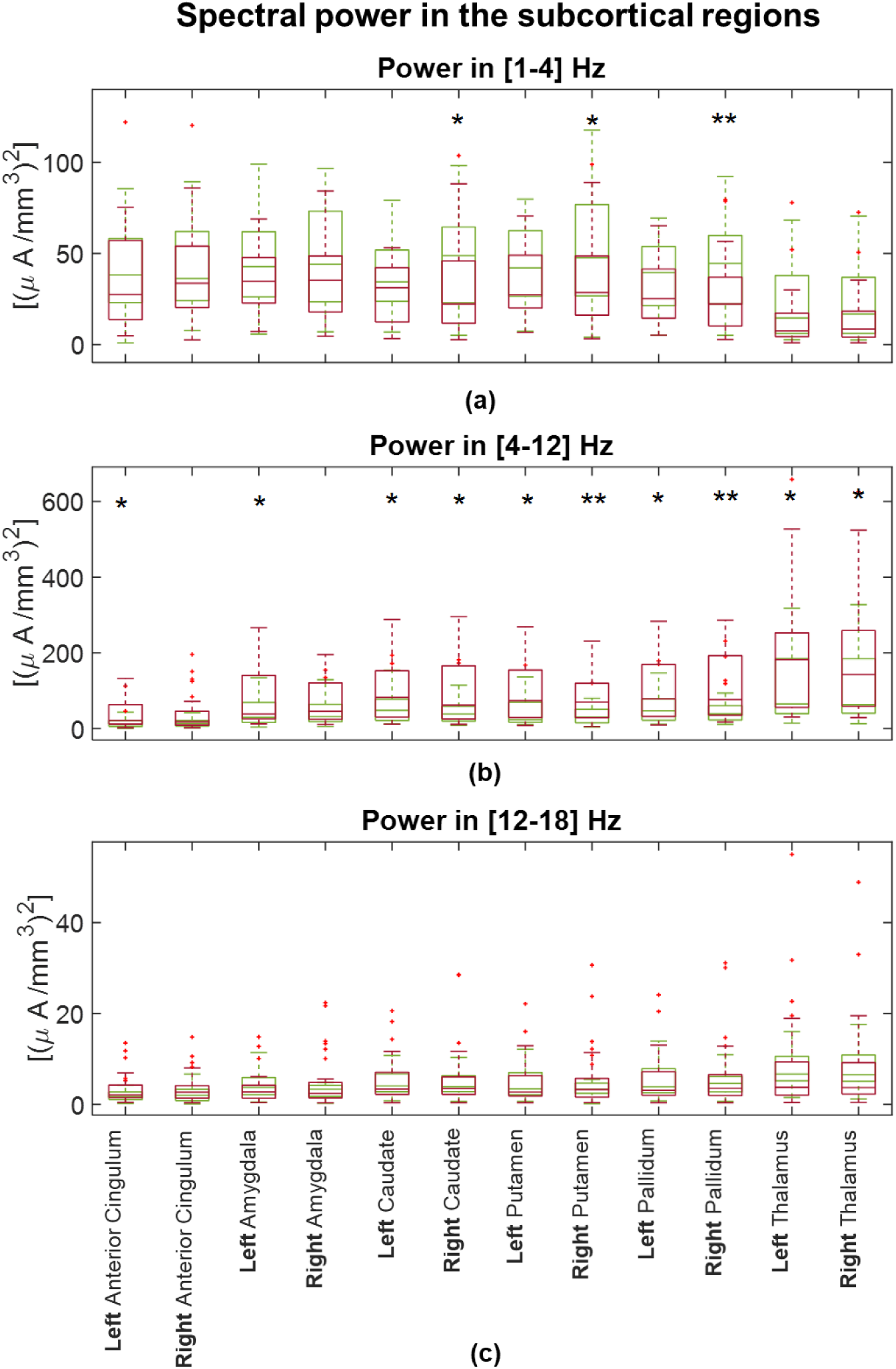
Boxplots to graphical illustrate the distribution of power of controls (green boxes) and patients (red boxes) in (a) [1-4] Hz, (b) [4-12] Hz and (c) [12-18] Hz. One star (*) stands for significant statistical difference with p<0.05 and two stars (**) for p<0.001. Power in [4-12] Hz significantly increases in patients compared to controls in all examined anatomical brain structures.

We found a significantly decreased power in delta [1-4] Hz and beta [12-18] Hz frequency bands in patients compared to controls in all investigated ROIs, when evaluating the results at the *population* level (Figure 1). At the *single-subject* level, delta power was significantly decreased in patients vs controls in the right caudate, putamen, and pallidum (Figure 2a). There was no significant difference in beta power between the two groups in any investigated ROI at the *single-subject* level (see Figure 2c).

### Network metrics

The connectivity network measures that we performed in the [4-12] Hz frequency range, showed increased values in patients compared to controls at both levels. At the *population* level, the local efficiency measured in patients was higher than in controls in all examined subcortical ROIs (see Figure 3). At the *single-subject* level, the global efficiency was significantly higher (p<0.05) in patients (mean ± standard deviation: 0.0129 ±0.0021) than in controls (mean ± standard deviation: 0.0126±0.0019). Considering all brain regions, the local efficiency tended to be higher in patients compared to controls (see Supplementary Fig. S2 online) but the significant differences corresponded only to the right precentral, amygdala and caudate regions (p<0.05). We observed significant correlations between the local efficiency and power in the [4-12] Hz frequency range in subcortical ROIs but it was not generalized among all twelve subcortical ROIs (see Supplementary Fig. S3 online). No significant correlations were found between the local efficiency and power in delta and beta bands. All the network measures computed on the twelve selected ROIs showed significantly higher values in patients than in controls in the right amygdala. The strength, local efficiency, and clustering coefficient of the right caudate were significantly higher in patients than controls, while there was no significant difference between the groups in the outflow from this ROI. There were no significant differences in any network metric in the anterior cingulate, thalamus, pallidum, or putamen (see Figure 4).

**Figure 3.**
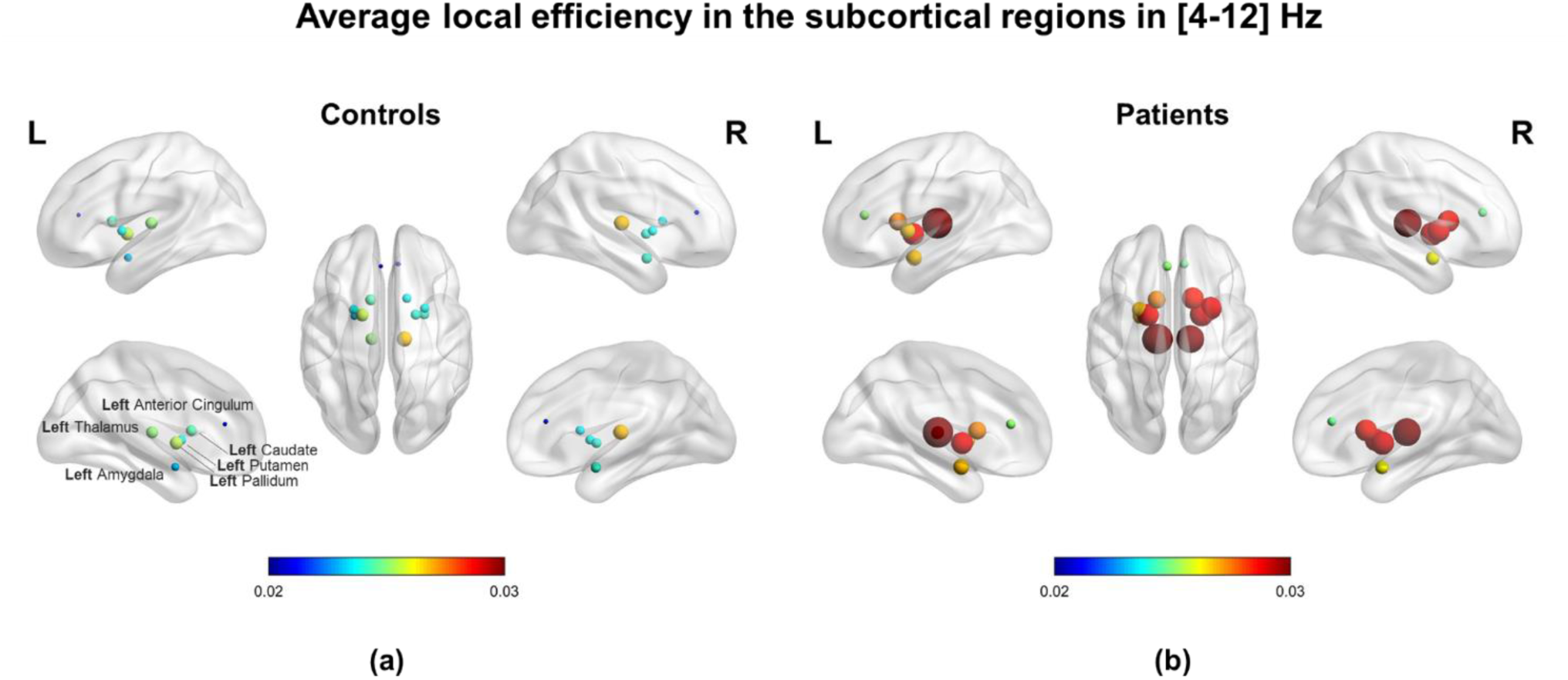
Local efficiency computed in the two *population subjects* representing (a) controls and (b) patients. Note that all subcortical regions of interest (ROIs) revealed higher values for patients than controls corresponding to the same tendency observed in all ROIs of the brain at the *single-subject* level (see Supplementary Fig. S1 – S2 online). The efficiency for each ROI is represented by a sphere centered on the cortical region, whose radius is linearly related to the magnitude. Such information is also coded through a color scale.

**Figure 4.**
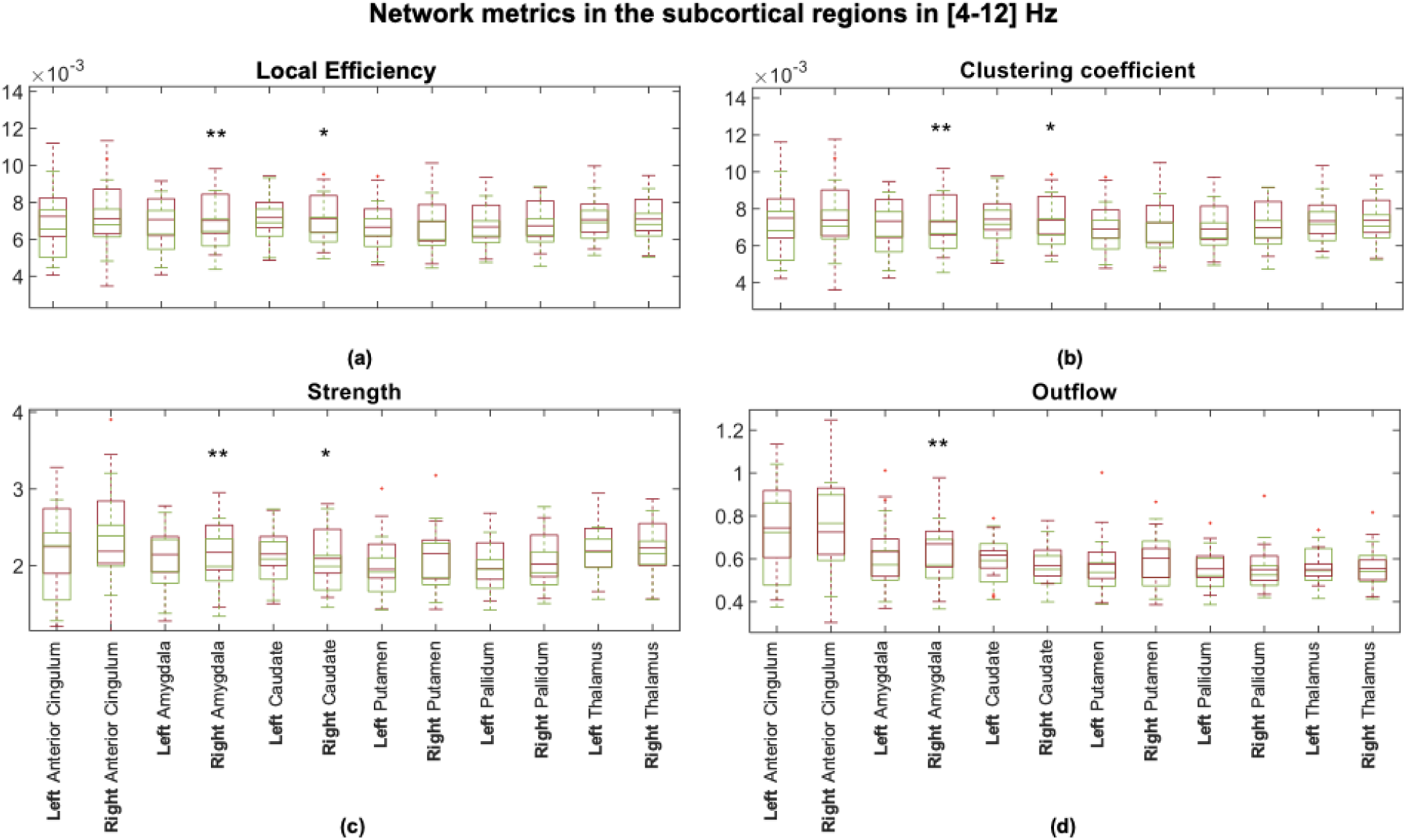
Boxplots to graphically illustrate the distribution of (a) local efficiency, (b) clustering coefficient, (c) strength, and (d) outflow in controls (green boxes) and patients (red boxes). One star (*) stands for significant statistical difference with p<0.05 and two stars (**) for p<0.001. All network metrics that refer to the right amygdala significantly differ between controls and patients (p<0.001), applying the Bonferroni correction (p<0.05/12 → p<0.0042).

There were no statistical differences in the network metrics estimated between the left and right hemisphere in each subject. The laterality indices showed that neither controls, nor patients had a lateralization in connectivity results of the six investigated deep brain structures. No significant differences in the laterality indices were observed comparing controls and patients.

### Effect of medication on network impairments

We found no correlation of the connectivity results with the intake of benzodiazepines, while there was a significant relationship between the global efficiency as predictor of the intake of AD/AP/MS medication (AD/AP/MS ∼ 1 + GE + GE^2^; Root Mean Squared Error: 0.716; *R*^2^ = 0.59; F-statistic vs. constant model: 18.7, p = 1.52e − 05). The global efficiency decreased with increasing medication score (see Figure 5). We observed no significant correlation (*R*^2^ < 0.05 and p > 0.8) between the connectivity results and any of the parameters that describe the status of depression (MADRS score, CGI score, illness duration, and the number of episodes) or the demographic profile (age and education level).

**Figure 5.**
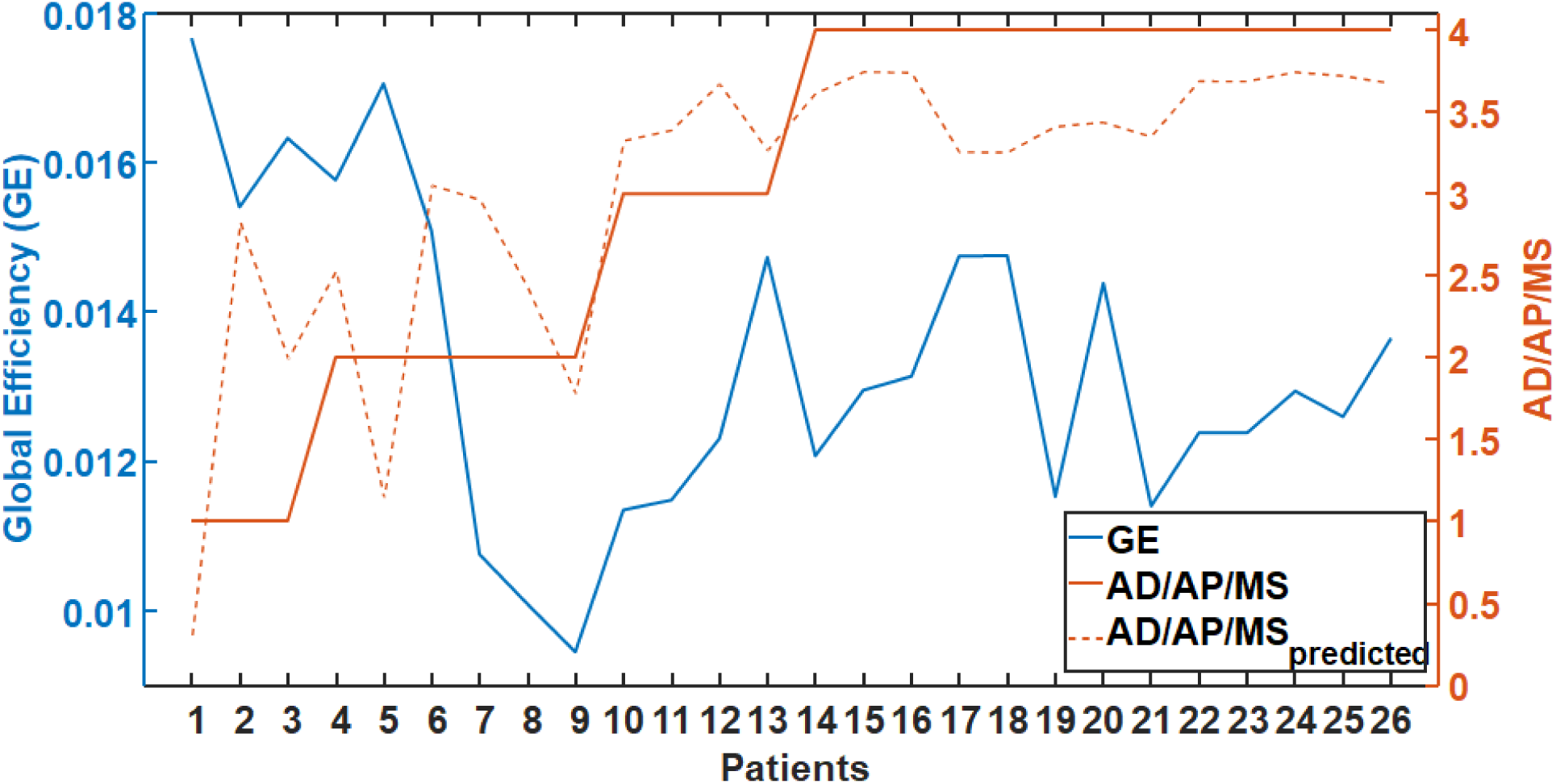
Relationship between the intake of antidepressants/antipsychotics/mood stabilizers (AD/AP/MS) and the global efficiency (GE). Note that higher medication intake is associated with lower GE. The orange dotted line stands for the predicted value of AD/AP/MS for each patient using GE as predictor. For values of the AD/AP/MS medication scale the reader is referred to the legend of Table 2.

## Discussion

In this study, we investigated resting-state network alterations using iPDC on source signals of high-density EEG in patients with depression compared to healthy controls. We explored the directed functional connectivity of the amygdala, anterior cingulate, putamen, pallidum, caudate, and thalamus, among them and with all the other brain regions in the time and frequency domain. We exploited the Kalman filter algorithm ^67^ assuming that resting state EEG segments were multiple realizations of the same process. Although we collapsed the temporal dimension to evaluate the network metrics, we decided to use a time-varying adaptive algorithm instead of a stationary autoregressive model to take into account the possible non-stationarity of the EEG signal and to more accurately capture this variability before collapsing the time with a summary measure, e.g., the median.

**Table 1.** Demographic data.

| Characteristic | Patients<br>(n = 26) | Controls<br>(n = 25) | *t*-value | df | p-<br>value |
| --- | --- | --- | --- | --- | --- |
| Age: mean ± SD | 51.9 ± 9.1 | 49.5 ± 8.7 | 0.97 | 49 | 0.34 |
| Gender: female, *n* | 11 | 10 |  |  |  |
| Education<sup>a</sup>: mean ± SD | 1.9 ± 0.9 | 2.3 ± 0.7 | -1.70 | 49 | 0.10 |

**Table 2.**
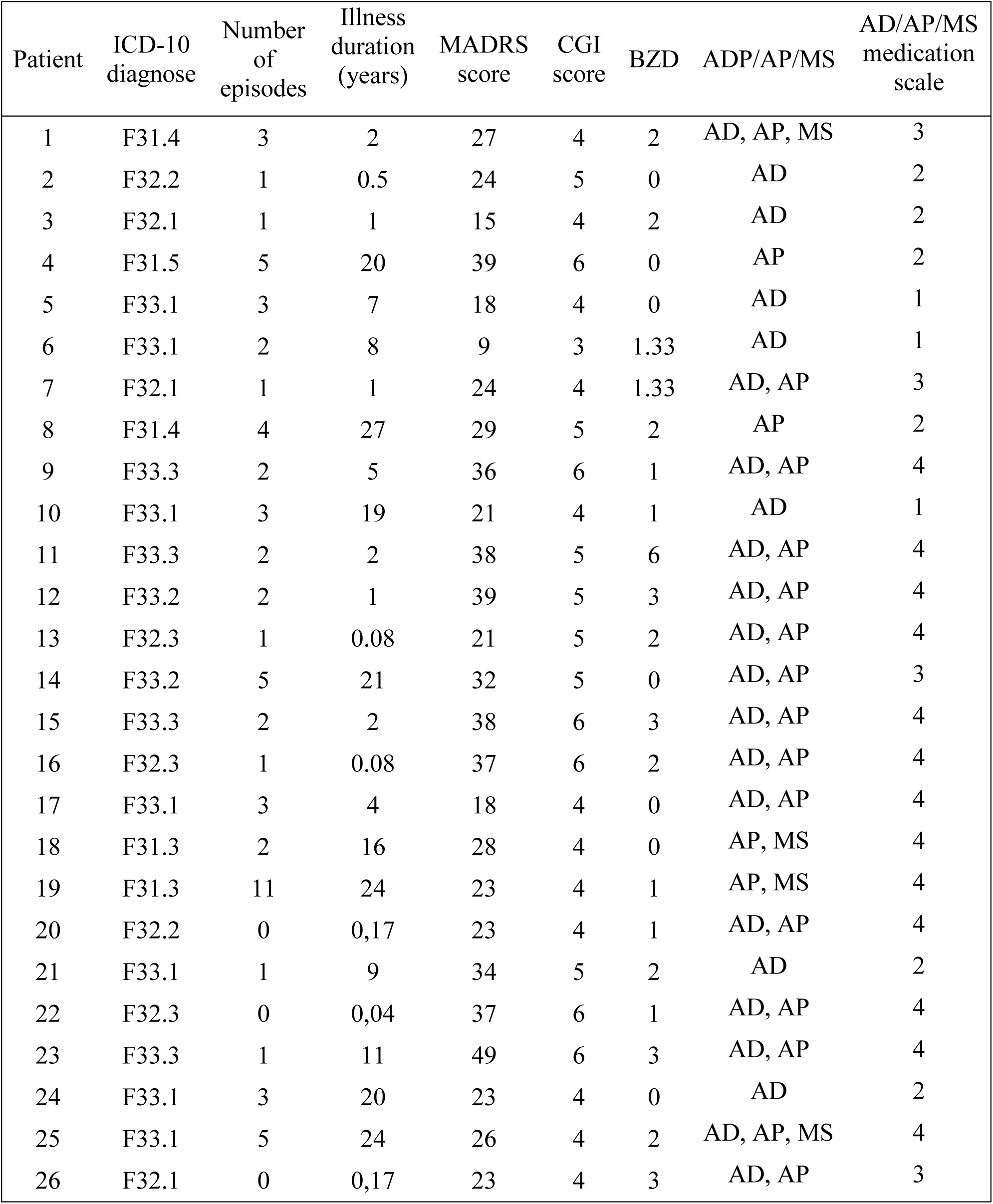
Patient characteristics.

To sum up, we demonstrated that in patients with moderate to severe depression: (1) the directed functional connectivity was significantly increased compared to controls in the right amygdala and the right caudate; (2) the power in theta and alpha frequency bands was significantly increased compared to controls in all investigated brain anatomical structures; (3) higher medication intake was associated with lower overall driving from the investigated regions.

### Increased right amygdala directed functional connectivity in depression

The most robust finding in our study was an abnormally increased directed functional connectivity in the right amygdala during resting-state in depressive patients. Even though the left-right asymmetry was not demonstrated by the laterality indices, a right-lateralized hyper-connectivity, as revealed with all the computed network metrics, was observed in the amygdala. We observed an increase in outgoing connections from the right amygdala as reflected with significantly higher outflow and strength in patients compared to controls. Moreover, we found a hyper-connectivity in the local networks of the right amygdala as reflected with significantly higher local efficiency and clustering coefficient in patients compared to controls.

We also found a significantly higher global efficiency in patients compared to healthy controls. This network feature had the same trend at the population level. Namely, we observed abnormally increased local efficiency of all examined deep brain structures in depressive patients. The efficiency measures the ability of a neural network to integrate and combine information. The deeper regions have a key role as hubs of the large-scale brain networks, so changes in their local connectivity properties might have also led to connectivity changes in the whole brain.

The amygdala is involved in processing salient stimuli ^68 69^ and has been implicated as one of the central hubs within the affective salience network ^70 71 72^. There is converging evidence from the neuroimaging studies that points to an abnormally increased connectivity and heightened activation of the amygdala in major depressive disorder (MDD) patients ^73 74^, Reduced connectivity ^75 76^ and anomalous subregional functional resting-state connectivity of the amygdala ^77^ were also reported. Distinct network dysfunctions in MDD were suggested to underlie adult-specific amygdala resting-state fMRI connectivity impairment within the affective network, presumably reflecting an emotional dysregulation in MDD ^76^. Hyperconnectivity between the amygdala, default mode network and salience network was also found to be related to depressive symptoms suggesting to underlie the poststroke depression in temporal lobe lesions ^78^. Unfortunately, the directionality of connections, which might be of interest when considering a structure as a potential DBS-target for treatment of TRD, cannot be inferred from these functional studies. There are only rare EEG-based connectivity studies focusing on depressive symptoms ^58 59 60 79^ that are, however, conducted only on a non-clinical population ^79^ or with connectivity parameters calculated at the sensor level ^57 58 59 60 61^. Authors of one of these studies ^79^ suggested an inability of the left dorsolateral prefrontal cortex to modulate the activation of the left temporal lobe structures to be a crucial condition for ruminative tendencies. Interestingly, in the current study we demonstrated an abnormal increase in directed functional connectivity arising from the right amygdala. This increased connectivity in depressive patients could reflect an abnormal functioning of the right amygdala. Such dysfunction might represent an impaired bottom-up signaling for top-down cortical modulation of limbic regions, leading to an abnormal affect regulation in depressive patients.

The increased functional connectivity in amygdala is likely related to structural changes observed in depression. Enlarged amygdala volumes was found in first-episode depressive patients that positively correlated with severity of depression ^80^. Higher grey matter volume was detected in bilateral amygdala of TRD patients compared to non-TRD patients, irrespective whether the patients presented bipolar or unipolar features and was suggested to reflect vulnerability to chronicity, revealed by medication resistance ^81^. Larger right amygdala volume was, however, also suggested to be associated with greater chances of remission in bipolar disorder ^82^.

In our study we aimed to investigate the directed functional connectivity in amygdala to provide knowledge on neurobiology of depression that is needed to evaluate this structure as a possible candidate for DBS treatment in depression. Despite myriad of DBS targets for treating depression tested in humans ^20^, the amygdala is not among them. The possible safety and utility of DBS in the amygdala could only be inferred from studies, in which the amygdala-DBS was performed for other neuropsychiatric diagnoses, such as epilepsy ^83 84 85 86^, post-traumatic stress disorder ^87 88^, and autism ^89^. In one of these studies transient stimulation-related positive shift in mood was observed ^84^. Particularly, the stimulation of the right amygdala induced a transient decrease in the negative affective bias, i.e. the tendency to interpret ambiguous or positive events as relatively negative. In this case study, an epileptic patient with MDD rated the emotional facial expressions as more positive with stimulation than without. The stimulation effect might have been associated with a transient normalization of likely impaired function of the right amygdala in that patient. We can only speculate, whether this dysfunction was in terms of hyper-connectivity similar to that observed in our study and whether it was temporally decreased by inhibitory effect of the stimulation.

### Increased right caudate directed functional connectivity in depression

We demonstrated that during resting state, patients had significantly higher right caudate directed functional connectivity than healthy controls. Despite no significant difference between groups in the caudate outflow, we observed an abnormally increased strength of outgoing connections from the right caudate in patients. Moreover, we found a hyper-connectivity in the local networks of the right caudate as reflected with significantly higher local efficiency and clustering coefficient in patients compared to controls. Caudate hyperactivation and increased caudate-amygdala and caudate-hippocampus fMRI connectivity during stress was previously reported in remitted individuals with recurrent depression ^90^. The here observed EEG-based functional caudate hyperconnectivity suggests striatal dysfunction even during resting-state in depressed patients. Our finding is consistent with a compelling evidence directly associating cortico-basal ganglia functional abnormalities with primary bipolar and unipolar spectrum disorders ^91^. Deficits in resting-state default-mode network connectivity with the bilateral caudate were suggested to be an early manifestation of MDD ^92^. Reduced grey matter volume in the bilateral caudate ^93 94 95 12^, diffusion tensor imaging-based hypoconnectivity between the right caudate and middle frontal gyrus ^96^, and altered functional connectivity of the right caudate with the frontal regions ^94^ was observed in MDD patients. In a post-mortem morphometric study in late-life depressive subjects, reduction in neuronal density was found in both the dorsolateral and ventromedial areas of the caudate nucleus ^97^. Associations between increased white matter lesion volumes and a decreased right caudate volume in the late-life depression was reported ^98^. In mild to moderately depressed patients no change in caudate gray matter volumes were found ^99^ suggesting inverse correlation between the caudate volume and severity of depression.

We found no significant differences in any network metric in the putamen, pallidum, thalamus, and anterior cingulate. It is possible, however, that examining these structures as a whole might be insensitive to different changes in their relevant subregions. Only the medial part of the thalamus is expected to play a role in the experience of affect ^73 100^. Reduced activity in the dorsal ACC but increased activity in the subgenual ACC have been found in acute depression in functional imaging studies ^101 102^. Moreover, we must take into account the limitations of our methodological approach, i.e. the source localization of the EEG activity in the subcortical regions. We have to keep in mind that the spatial resolution in detecting and distinguishing neighboring brain regions is about 24 mm ^103^. Therefore, our results in the caudate, putamen and pallidum are probably overlapping due to smearing of the sources. Keeping in mind these limitations and with respect to the lower robustness of our findings in the caudate, we can just encourage researchers to further investigate the neuropathophysiology of depression associated with the caudate nucleus functioning. More evidence from neuroimaging studies is needed to provide arguments for the next caudate-DBS tests in treating TRD. In an early case study, DBS of the ventral caudate nucleus markedly improved symptoms of depression in a patient with MDD and comorbid obsessive-compulsive disorder ^104^. No change in depressive symptoms, however, was recently observed during the stimulation of the caudate in a study of three TRD patients ^105^ and authors concluded the caudate to be less promising DBS target than the nucleus accumbens.

### Increased theta and alpha powers in depression

We found a significantly higher power in the theta and alpha frequency bands in the depressed compared to the healthy control group in all the investigated subcortical structures consistently at both the population and single-subject levels. The power decrease in the beta and delta frequency bands was observed only in the right striatum at both levels.

Our findings might be in line with previous observations in the sensor space of the scalp EEG. Abnormally high power in alpha ^106 107 108^ and theta ^106 109 108^ frequency bands in parietal and occipital regions were found in depressed patients, lower than normal beta and delta power were also reported ^108^. Recent evidence points, however, to opposite power changes showing that theta and alpha power might decrease, while beta power increases in depression ^110^. Moreover, the same study reported negative association of the posterior alpha power with the depression severity. While changes in cortical theta and alpha activity were suggested to be inversely related to the level of cortical activation, enhancement of the cortical beta power was suggested to reflect higher level of anxiety symptoms in depressed patients ^106^. To the best of our knowledge there is only one study that directly recorded electrophysiological activity in subcortical structures in depressive patients. In this study, a larger alpha activity in MDD patients compared with obsessive compulsive disorder was found in the limbic DBS targets (the anterior cingulate and the bed nucleus of the stria terminalis) ^111^. Moreover, in the same study, the increased alpha power correlated with severity of depressive symptoms. Nevertheless, in spite of parallels with prior reports, the current link between the power changes in subcortical structures and depression awaits replication.

### Lower network impairments with more medication

We found an inverse relationship between the intake of medication and the impairment of the investigated networks. Particularly, increased intake of antidepressants, antipsychotics, and mood stabilizers was associated with reduction of the global efficiency. This finding might be related to the pharmacological effect on the brain activity, i.e. a change towards the normalization of the hyper-connectivity in the cortico-striatal-pallidal-thalamic and limbic networks. The low sample size and great variability in medication made it, however, impossible to examine any potential influence of medication on the network impairments by comparing patients receiving a specific drug with those not receiving it. To summarize the various medications, an ordinal variable was used that is only a rough measurement of medication usage. Moreover, the duration of the illness rather than the duration of the specific drug intake was considered in our study. Only doses of medication actually taken at the time of experiment were taken into consideration. The possible accumulated effect of specific drugs on connectivity results, thus, cannot be assessed. Therefore, the observed relationship between the global efficiency and medication should be viewed with caution. Interestingly, we have not found significant correlation between the global efficiency and intake of benzodiazepines. This negative finding suggests that even though benzodiazepines are known to have an effect on electrophysiological correlates of brain functions, the network properties might not be influenced. There were no significant correlations between the connectivity results and depressive symptom severity or other parameters describing the status of depression within the patient group. We suppose that heterogeneity of our dataset, in which patients with different disorders were included, might underlie this observation. We also found no relation between the connectivity results and education level or age. This finding suggests independence of the observed impairment on these demographic variables, however, the current sample size might be insufficient for such investigations.

### Limitations of the study

We here report sources of scalp-recorded electrophysiological brain activity in deep brain structures. We are aware of the limitations of EEG in sensing deep brain structures. However, previous work using simulations and source reconstruction provided indirect evidence for the detectability of subcortical sources in non-invasive EEG and magnetoencephalographic recordings ^112 113 114 115^. Moreover, recent simultaneous scalp and intracranial recordings directly demonstrated that activity in deep brain structures spread to the scalp ^103 116^. While Seeber and colleagues ^103^ used individual head models that improve source localization precision, a generic head model was used in the magnetoencephalographic study by Pizzo et al. ^116^, similar to the approach used in our study. Nevertheless, the results that we report have to be interpreted with caution and need further validation by intracranial recordings in future studies.

### Conclusions

We found an overall increase in power in theta and alpha frequency bands in depressive patients compared to healthy controls in the subcortical regions constituting the cortico-striatal-pallidal-thalamic and limbic circuits. The network measures showed a higher than normal functional connectivity arising from the right amygdala in depressive patients. The amygdala seems to play an important role in neurobiology of depression. Resting-state EEG directed functional connectivity is a useful tool for studying abnormal brain activity in depression.

## Methods

### Subjects

Data were collected from 26 depressive patients and 25 healthy controls. The two groups were matched by gender and there were no significant differences in age or education (see Table 1). On a subsample of this dataset we recently showed that the severity of depressive symptoms correlates with resting-state microstate dynamics^117^. The patients were recruited at the Department of Psychiatry, Faculty of Medicine, Masaryk University and University Hospital Brno, Czech Republic. The diagnostic process had two steps and was determined based on the clinical evaluation by two board-certified psychiatrists. First, the diagnosis was made according to the criteria for research of the International Classification of Disorders (ICD-10). Second, the diagnosis was confirmed by the Mini International Neuropsychiatric interview (M.I.N.I.) according to the Diagnostic and Statistical Manual (DSM-V). All patients were examined in the shortest time period after the admission and before the stabilization of treatment, typically during their first week of hospitalization. All patients met the criteria for at least a moderate degree of depression within the following affective disorders: bipolar affective disorder (F31), depressive episode (F32), recurrent depressive disorder (F33). Exclusion criteria for patients were any psychiatric or neurological comorbidity, IQ < 70, organic disorder with influence on the brain function, alcohol dependence or other substance dependence. All patients were in the on-medication state with marked interindividual variability in specific medicaments received. Control subjects were recruited by general practitioners from their database of clients. Control subjects underwent the M.I.N.I. by board-certified psychiatrists, to ensure that they had no previous or current psychiatric disorder according to the DSM-V criteria. The scores on the Montgomery-Åsberg Depression Rating Scale (MADRS), a specific questionnaire validated for patients with mood disorders ^118^ and the Clinical Global Impression (CGI) ^119^, a general test validated for mental disorders, were used to evaluate the severity of depressive symptoms in patients. The status of depression was further described with life time count of depressive episodes and illness duration. Medication in 24 hours preceding the EEG examination was also recorded (see Table 2). This study was carried out in accordance with the recommendations of Ethics Committee of University Hospital Brno with written informed consent from all subjects.

### EEG - data acquisition and pre-processing steps

Subjects were sitting in a comfortable upright position in an electrically shielded room with dimmed light. They were instructed to stay as calm as possible, to keep their eyes closed and to relax for 15 minutes. They were asked to stay awake. All participants were monitored by the cameras and in the event of signs of nodding off or EEG signs of drowsiness detected by visual inspection, the recording was stopped. The EEG was recorded with a high density 128-channel system (EGI System 400; Electrical Geodesic Inc., OR, USA), f_s_ = 1kHz, and Cz as acquisition reference.

Five minutes of EEG data were selected and visually assessed. Noisy channels with abundant artifacts were identified. EEG signal was band-pass filtered between 1 and 40 Hz with a 2nd-order Butterworth filter avoiding phase-distortion. Subsequently, in order to remove physiological artifacts, e.g. ballistocardiogram and oculo-motor artifacts, infomax-based Independent Component Analysis ^120^ was applied on all but one or two noisy channels. Only components related to ballistocardiogram, saccadic eye movements, and eye blinking were removed based on the waveform, topography and time course of the component. Then, the cleaned EEG recording was down-sampled at f_s_ = 250 Hz and the previously identified noisy channels were interpolated using a three-dimensional spherical spline ^121^, and re-referenced to the average reference. For the following analyses, thirty 2-s EEG epochs free of artifacts were selected per subject. All the pre-processing steps were done using the freely available Cartool Software 3.70, programmed by Denis Brunet ^122^ and custom functions in MATLAB® R2018b.

### EEG source estimation

We applied the LAURA algorithm implemented in Cartool ^122^ to compute the source reconstruction taking into account the patient’s age to calibrate the skull conductivity ^123 124 125^. The method restricts the solution space to the gray matter of the brain. Then, the cortex was parcellated into the 90 Automated Anatomical Labeling brain regions ^126^. The dipoles in each ROI were represented with one unique time-series by a singular-value decomposition ^127^.

### Time-variant multivariate autoregressive modeling

The cortical waveforms computed after applying the singular-value decomposition, were fitted against a time-variant (tv) multivariate (MV) autoregressive (AR) model to overcome the problem of non-stationarity of the EEG data. If the EEG data are available as several trials of the same length, the cortical waveforms computed from the EEG data generates a collection of realizations of a multivariate stochastic process which can be combined in a multivariate, multi-trial time series ^127 128 67^. The tv-MVAR matrices containing the model coefficients were computed in the framework of a MATLAB toolbox (code available upon reasonable request to the authors) that implements the adaptive Kalman filtering and information Partial Directed Coherence (iPDC) in the source space ^67 129 130^. The model order of the tv-MAR and the Kalman filter adaptation constant were chosen applying the method proposed by Rubega and colleagues ^128^, i.e., evaluating the partial derivatives of a residual minimization function obtained varying simultaneously both p (p € [1, 15]) and c (c € [0, 0.03]). By means of the model coefficients, we computed the parametric spectral power density and the iPDC absolute values for each subject. For each patient, we obtained a 4-dimensional matrix [ROIs x ROIs x frequency x time] that represented the directed information flow from one ROI to another for each frequency at each time sample. In this way we performed the analysis on the *single-subject* level to compare the two groups quantitatively.

Since the features in the power spectra were consistent among subjects in the same population (patients vs controls), we also performed the analysis on the *population* level. A *population subject* was built by estimating the tv-MVAR model, where each trial in the input was a different subject. One power spectral density matrix and one connectivity matrix [ROIs x ROIs x frequency x time] were obtained for each group (controls and patients). In other words, subjects were combined as trials, assuming respectively humans as multiple realizations of their own brain processes, with the purpose to show that the two approaches, i.e., *single subject* and *population*, give equivalent results in differentiating patients vs controls. In the last decade, population-based approaches were successfully exploited in computer simulations engineered to evaluate the safety and limitations of closed-loop control treatment algorithms ^131 132^. Population-based approaches for MVAR/PDC modelling are currently lacking and this might be considered a first attempt justified by the consistent features estimated in the frequency domain among subjects belonging to the same population (patients vs controls). Further details on the connectivity estimation are reported in the Supplementary Information.

### Network metrics

In order to study the peculiarities of the brain network in patients vs controls, the brain was represented as a digraph defined by a collection of nodes and directed links (directional edges). Nodes in the brain network represent brain regions, i.e., the 90 ROIs, while the directed links represent the values computed by iPDC. Thus, the weight of such link can vary in the interval [0-1] and it represents the amount of mutual information flowing between ROIs. We defined twelve ROIs, including the bilateral amygdala, anterior cingulum, thalamus, putamen, caudate, and pallidum, to examine the directed functional connectivity between these seeds and the whole brain. Significant differences in power between patients and controls were observed in the *single-subject* level in alpha and theta frequency bands in all these six anatomical structures. Therefore, we restricted the network analysis to this [4-12] Hz frequency range. To evaluate how much the system is fault tolerant and how much the communication is efficient, the global efficiency for the whole brain and the local efficiency, clustering coefficient, strength and outflow for each of these twelve investigated ROIs were computed. To compute all the graph measures, the scripts and functions implemented on the freely available MATLAB toolbox ^133^ were customized.

### Global efficiency

Global efficiency is defined as the average minimum path length between two nodes in the network. This measure is inversely related to topological distance between nodes and is typically interpreted as a measure of the capacity for parallel information transfer and integrated processing ^134^.

### Local efficiency

Local efficiency is defined as the average efficiency of the local subgraphs ^135^, i.e. the global efficiency computed on the neighborhood of the node. It reflects the ability of a network to transmit information at the local level. This quantity plays a role similar to the clustering coefficient since it reveals how much the system is fault tolerant, i.e., it shows how efficient the communication is between the first neighbors of *i* when *i* is removed.

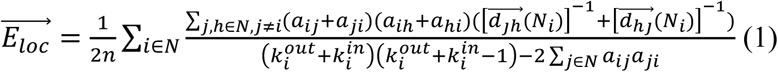

where 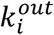 is the out-degree of node *i*, 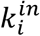 is the in-degree of node *i*, and *a*_*ij*_ is the connection status between node *i* and node *j*, i.e., *a*_*ij*_ = 1 if the link between *i* and *j* exists, *a*_*ij*_ = 0 otherwise. *N* is the set of nodes in the network. *n* is the number of nodes and 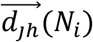 is the length of the shortest directed path between *j* (any node in the network) and *h* (any node that neighbors with *i*).

### Clustering coefficient

Clustering coefficient reflects the prevalence of clustered connectivity around an individual brain region ^136^:

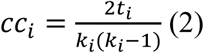

where *t*_*i*_ are the number of triangles around the node *i*, and *k*_*i*_ is the degree of node *i*, i.e., the number of links connected to node *i*. In our case of a weighted directed network, a weighted directed version of clustering coefficient was used ^137^:

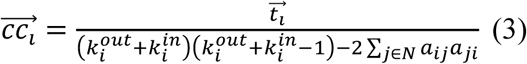

where 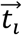 are the number of directed triangles around the node *i*, 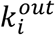 is the out-degree of node *i*, 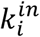 is the in-degree of node *i*, and *a*_*ij*_ is the connection status between the nodes *i* and *j*, i.e., *a*_*ij*_ = 1 if the link between *i* and *j* exists, *a*_*ij*_ = 0 otherwise. *N* is the set of nodes in the network.

### Strength and outflow

Finally, the connectivity patterns between the different cortical regions were summarized by representing the strength that quantifies for each node the sum of weights of all links connected to the node and the total outflow from a region toward the others, generated by the sum of all the statistically significant links obtained by application of the iPDC. The greatest amount of information outflow depicts the ROI as one of the main sources (drivers) of functional connections to the other ROIs ^138^.

### Laterality

For all the network metrics explained in the previous paragraph, we also computed a laterality index, which is defined as 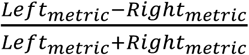 to test if the measures significantly differentiate between the two hemispheres. Laterality index and all network metrics were calculated for both groups.

### Statistical analysis

To assess whether or not the changes in the network metrics were statistically significant between patients and controls, paired Student’s t-tests were computed under the hypothesis of normal distribution of samples (Lilliefors test), otherwise Wilcoxon rank-sign tests were considered. To test whether the age and education level predict the values of the spectral power distribution and the network metrics in patients, a multiple linear regression was performed. We also tested the influence of the clinical data on the connectivity results. A multiple linear regression was performed exploiting correlation of the connectivity results with four variables describing the status of depression and two variables describing the medication status in terms of the intake of benzodiazepines (BZP), antidepressants, antipsychotics, and mood stabilizers (AD/AP/MS). These six clinical variables are provided for each patient in Table 2. We checked through the following multiple linear regression models (4) (5), if the response variable Y depends on a number of predictor variables *X*_*i*_:

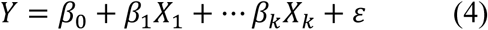

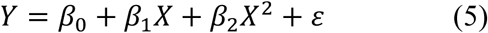

where the *ε* are the residual terms of the model and β_0_, β_1_, β_2_, …, β_k_ are the *k* regression coefficients. Both the clinical data and the power and network metrics were used once as predictors and once as response variables.

### Ethics statement

All participants gave their written informed consent prior to the experiment and the study received the approval of the Ethics Committee of University Hospital Brno in Brno, Czech Republic. All experiments of this study were performed in accordance with relevant guidelines and regulations.

## Acknowledgments

This study was supported by the European Union Horizon 2020 research and innovation program under the Marie Sklodowska-Curie grant agreement No. 739939, by Ministry of Health, Czech Republic - conceptual development of research organization (University Hospital Brno - FNBr, 65269705), by the Swiss National Science Foundation (grant No. 320030_184677), and by the National Centre of Competence in Research (NCCR) “SYNAPSY–The Synaptic Basis of Mental Diseases” (NCCR Synapsy Grant # “51NF40 – 185897). CMM and MR were supported by the Swiss National Science Foundation (Sinergia project CRSII5_170873). The funding sources had no role in the design, collection, analysis, or interpretation of the study. The authors wish to thank Martin Seeber, Patrik Wahlberg, David Pascucci, and Gijs Plomp for providing useful comments on the manuscript.

## Author Contributions

AD – designed the study, performed the preprocessing, and wrote the initial draft; RB and JH – were responsible for patient recruitment and clinical assessment; EH – collected the EEG data; SF and ŠO – were involved in the clinical assessment; CMM – served as an advisor, MR – performed the analysis, wrote the initial draft, and was responsible for the overall oversight of the study. All authors revised the manuscript.

## Table legends

Table 1. ^a^Education was classified into three levels: 1 = no high school, 2 = high school, 3 = university studies

Table 2. F31.3 - Bipolar affective disorder, current episode mild or moderate depression; F31.4 - Bipolar affective disorder, current episode severe depression without psychotic symptoms; F31.5 - Bipolar affective disorder, current episode severe depression with psychotic symptoms; F32.1 - Moderate depressive episode; F32.2 - Severe depressive episode without psychotic symptoms; F32.3 - Severe depressive episode with psychotic symptoms; F33.1 - Recurrent depressive disorder, current episode moderate; F33.2 - Recurrent depressive disorder, current episode severe without psychotic symptoms; F33.3 - Recurrent depressive disorder, current episode severe with psychotic symptoms; BZD: benzodiazepine equivalent dose ^139^ AD - antidepressants (mirtazapine, citalopram, venlafaxine, vortioxetine, sertraline, trazodone); AP - antipsychotics (risperidone, olanzapine, quetiapine, amisulpride, aripiprazole); MS - mood stabilizers (valproate, lamotrigine, carbamazepine); AD/AP/MS medication scale: 1 – one medication in sub-therapeutic doses, 2 – one medication in therapeutic doses, 3 – combination of medications with one in therapeutic doses, 4 – combination of medications with more than one in therapeutic doses; MADRS (Montgomery–Åsberg Depression Rating Scale): score is between 0 and 60, the higher the score the higher the depressive symptom severity; CGI (Clinical Global Impression scale): healthy (1) – most extremely ill (7). Four patients were undergoing the first (patient 3) and second (patient 4 and 9) week of electroconvulsive therapy and the first week of repetitive transcranial magnetic stimulation (patient 5). No clinical effect of these neurostimulation treatments was apparent.

