## Supplemental Information for "Altered directed functional connectivity of the right amygdala in depression: high-density EEG study"

### Further insights on connectivity estimation

On one hand, modifications on the EEG power spectral density in the canonical rhythms (delta, theta, alpha, beta and gamma) permit to infer information on brain activity in different areas. On the other hand, mathematical models may help to elucidate the relationship among brain electric potentials allowing the quantitative analysis of the connections among the different brain regions.

Most of the studies have used approaches based on the definition of Partial Directed Coherence (PDC), i.e., a frequency domain representation of Granger-causality. The originally defined PDC is not scale invariant, i.e., arbitrary changes in the amplitudes of one time series can lead to substantial changes in PDC values. This deficiency was first addressed by introducing the notion of generalized PDC (gPDC) and then by introducing information PDC (iPDC). This weighted multivariate directed dependence measure aims to compute the relationship among partialized signals, inferring the mutual information rates between process portions[1]. The main feature is to adequately consider the prediction error covariance matrix in its formulation to weight the relationships among signals into correctly scaled information measurements, when stationarity zero-mean white noise processes also called innovation processes are either strongly correlated or when their variances differ considerably in size. $iPDC$ properly accounts for size effects in gauging connection strength, given its ability to express information flow in rigorous fashion, as reported in detail in[1 2].

In particular, $iPDC$ is a multivariate spectral measure to compute only the directed influences between any given pair of signals $(i, j)$ of a multivariate dataset. This information is condensed in a complex function $iPDC_{i \leftarrow j}(f)$ of the frequency $f$, which measures the relative interaction of the signal $j$ with regard to signal $i$ as compared to all $j$'s interactions to other signals in the multivariate dataset. While we refer the reader to[1] for further mathematical details, the procedure for computing $iPDC$ is briefly described by the following two steps.

In the first step, the cortical waveforms $\tilde{x}$ computed after applying the projection method described in [3], are fitted against a *time-variant* (tv) multivariate autoregressive (MVAR) model to overcome the problem of non-stationarity of the EEG data. When the EEG data are available as several trials of the same length, the cortical waveforms computed from the EEG data generates a collection of realizations of a multivariate stochastic process, which can be combined in a multivariate, multi-trial time series:

$$\widetilde{\boldsymbol{X}}(t) = \begin{bmatrix} \widetilde{\boldsymbol{x}}_1^{(1)}(t) & \cdots & \widetilde{\boldsymbol{x}}_d^{(1)}(t) \\ \vdots & \ddots & \vdots \\ \widetilde{\boldsymbol{x}}_1^{(K)}(t) & \cdots & \widetilde{\boldsymbol{x}}_d^{(K)}(t) \end{bmatrix} \quad t = t_1, .., t_N \qquad (1)$$

where $t$ refers to the time points, $N$ the length of the time-series, $K$ the number of trials and $d$ the number of ROIs.

Then the data in $\widetilde{\boldsymbol{X}}$ are fitted against a tvMVAR model in the general form:

$$\widetilde{\boldsymbol{X}}(t) = -\sum_{r=1}^{p} \boldsymbol{A_r}(t)\boldsymbol{X}(t-r) + \boldsymbol{W}(t) \qquad (2)$$

where $A_r(t)$ are the $[d\ x\ d]$ AR matrices containing the model coefficients, $W(t)$ is the innovation process with covariance matrix $\sum_w$, and $p$ is the model order, usually estimated by means of the Akaike Information Criteria for MVAR processes[3]. The General Linear Kalman filter approach is applied with the aim to estimate the coefficients of the time-variant AR matrices and the covariance matrix $\sum_w$ [4].

As the MVAR model is estimated, for each time-point $t$, having defined the complex matrix $\boldsymbol{B}(f)$ as:

$$\boldsymbol{B}(f) = \boldsymbol{I_d} - \sum_{r=1}^{p} \boldsymbol{A_r} e^{-j2\pi f} \qquad (3)$$

where $I_d$ is the identity matrix and $j$ is the imaginary unit in this equation, the $iPDC$ complex function from the time-series $j$ to the time-series $i$ is obtained by:

$$iPDC_{i\leftarrow j}(f) = \sigma_{w_{ii}}^{-1/2} \frac{b_{ij}(f)}{\sqrt{\boldsymbol{b_j^H}(f)\boldsymbol{\Sigma_w^{-1}} \boldsymbol{b_j}(f)}} \qquad (4)$$

where $\boldsymbol{b}_j\ (f)$ and $b_{ij}\ (f)$ are respectively the j-th column and the $(j, i)$-th element of matrix $\boldsymbol{B}(f)$, $\sigma_{w\_ii}$ is the $(i, i)$-th element of the innovation covariance matrix $\sum_w$, and the apex $H$ in $\boldsymbol{b}_j^H$ stands for Hermitian transpose, i.e., obtained from $\boldsymbol{b}_j$ by taking the transpose and then the complex conjugate of its components.

The complex function $iPDC_{i\leftarrow j}(f)$ of eq. (4) is usually analyzed in terms of its absolute value.

From the complex matrix computed in eq. (3), it is possible to estimate the parametric spectral power density (PSD):

$$PSD(f) = \boldsymbol{H}(f)\boldsymbol{\Sigma}_w\boldsymbol{H}(f)^T \qquad (5)$$

where $\boldsymbol{H}(f)$ is the signal transfer function equal to the inverse of $\boldsymbol{B}(f)$.

### Results on all regions of interest

Both the frequency analysis and the network analysis were computed for all source waveforms in the 90 regions of interest (ROIs) considered in the mathematical model for computing the iPDC matrix. The main assumption in the tv-MVAR model to compute the tv-iPDC is based on the hypothesis that all the source waveforms are included in the model itself. In the paper, the results reported are restricted to the pre-selection of twelve areas (the pre-selection is only in the visualization of the results, not in their computation) to answer to the main question of the work: Which structure within the cortico-striatal-pallidal-thalamic and limbic circuits reveals a disrupted resting-state directed functional connectivity? The twelve deep brain structures were selected to consider their potential implication in the deep brain stimulation treating treatment-resistant depression.

Here, the labels of the considered macroscopic brain structures are listed following the order of the AAL atlas: (1) left precentral gyrus; (2) right precentral gyrus; (3) left superior frontal gyrus; (4) right superior frontal gyrus; (5) left superior frontal gyrus, orbital part; (6) right superior frontal gyrus, orbital part; (7) left middle frontal gyrus; (8) right middle frontal gyrus; (9) left middle frontal gyrus, orbital part; (10) right middle frontal gyrus, orbital part; (11) left inferior frontal gyrus, pars opercularis; (12) right inferior frontal gyrus, pars opercularis; (13) left inferior frontal gyrus, pars triangularis; (14) right inferior frontal gyrus, pars triangularis; (15) left inferior frontal gyrus, pars orbitalis; (16) right inferior frontal gyrus, pars orbitalis; (17) left rolandic operculum; (18) right rolandic operculum; (19) left supplementary motor area; (20) right supplementary motor area; (21) left olfactory cortex; (22) right olfactory cortex; (23) left medial frontal gyrus; (24) right medial frontal gyrus; (25) left medial orbitofrontal cortex; (26) right medial orbitofrontal cortex; (27) left gyrus rectus; (28) right gyrus rectus; (29) left insula; (30) right insula; (31) left anterior cingulate gyrus; (32) right anterior cingulate gyrus; (33) left midcingulate area; (34) right midcingulate area; (35) left posterior cingulate gyrus; (36) right posterior cingulate gyrus; (37) left hippocampus; (38) right hippocampus; (39) left parahippocampal gyrus; (40) right parahippocampal gyrus; (41) left amygdala; (42) right amygdala; (43) left calcarine sulcus; (44) right calcarine sulcus; (45) left cuneus; (46) right cuneus; (47) left lingual gyrus; (48) right lingual gyrus; (49) left superior occipital gyrus; (50) right superior occipital gyrus; (51) left middle occipital gyrus; (52) right middle occipital gyrus; (53) left inferior occipital cortex; (54) right inferior occipital cortex; (55) left fusiform gyrus; (56) right fusiform gyrus; (57) left postcentral gyrus; (58) right postcentral gyrus; (59) left superior parietal lobule; (60) right superior parietal lobule; (61) left inferior parietal lobule; (62) right inferior parietal lobule; (63) left supramarginal gyrus; (64) right supramarginal gyrus; (65) left angular gyrus; (66) right angular gyrus; (67) left precuneus; (68) right precuneus; (69) left paracentral lobule; (70) right paracentral lobule; (71) left caudate nucleus; (72) right caudate nucleus; (73) left putamen; (74) right putamen; (75) left globus pallidus; (76) right globus pallidus; (77) left thalamus; (78) right thalamus; (79) left transverse temporal gyrus; (80) right transverse temporal gyrus; (81) left superior temporal gyrus; (82) right superior temporal gyrus; (83) left superior temporal pole; (84) right superior temporal pole; (85) left middle temporal gyrus; (86) right middle temporal gyrus; (87) left middle temporal pole; (88) right middle temporal pole; (89) left inferior temporal gyrus; (90) right inferior temporal gyrus.

From Supplementary Fig. S1, it is possible to qualitatively notice differences in the connectivity relationships among the 90 ROIs of the tv-MVAR model of Eq. 2. In order to quantitatively estimate the differences between controls and patients at the *population* level, graph metrics were extrapolated from this kind of matrices.

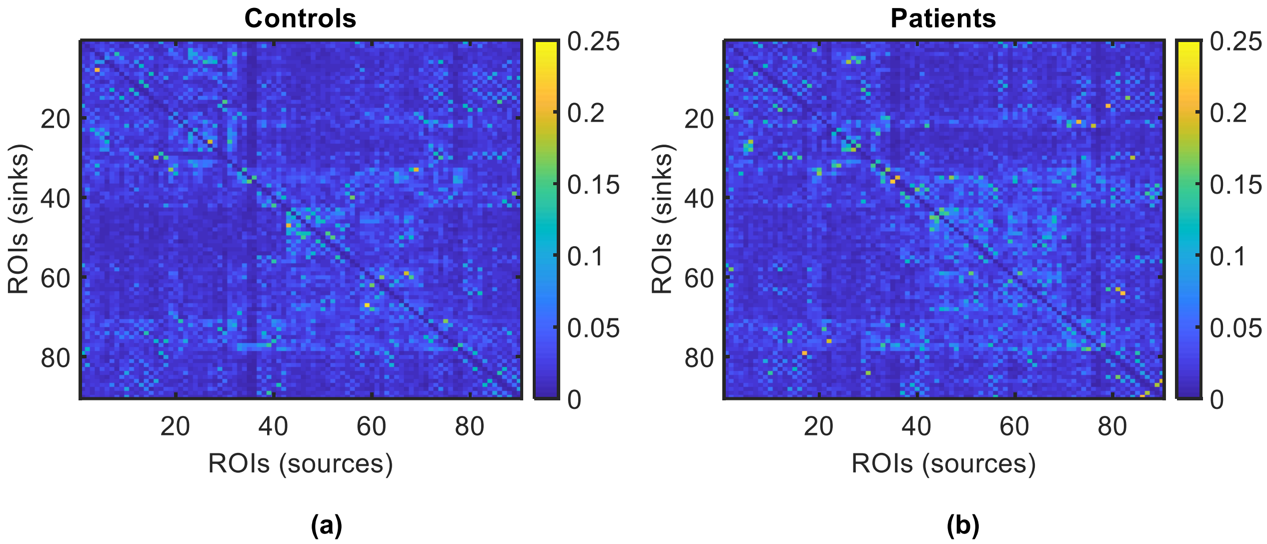


**Supplementary Fig. S1**: Absolute values of iPDC at the *population* level. Magnitude of iPDC values computed in two *population* subjects (i.e., each trial in input of the tv-MVAR model of Eq. 2 corresponds to the source waveforms computed from a different subject) representing (a) controls and (b) patients. The values reported were averaged over time and frequency. The ROIs are listed following the order of the AAL atlas. The direction of the reported information flow goes from the x-axis, i.e, ROIs (sources), to y-axis, i.e, the ROIs (sinks).

In Supplementary Fig. S2, we reported the values of the local efficiency for all the sources considered in the model at the *single-subject* level. Besides the right amygdala and caudate, also the right cuneus resulted significantly different between controls and patients (p<0.001).

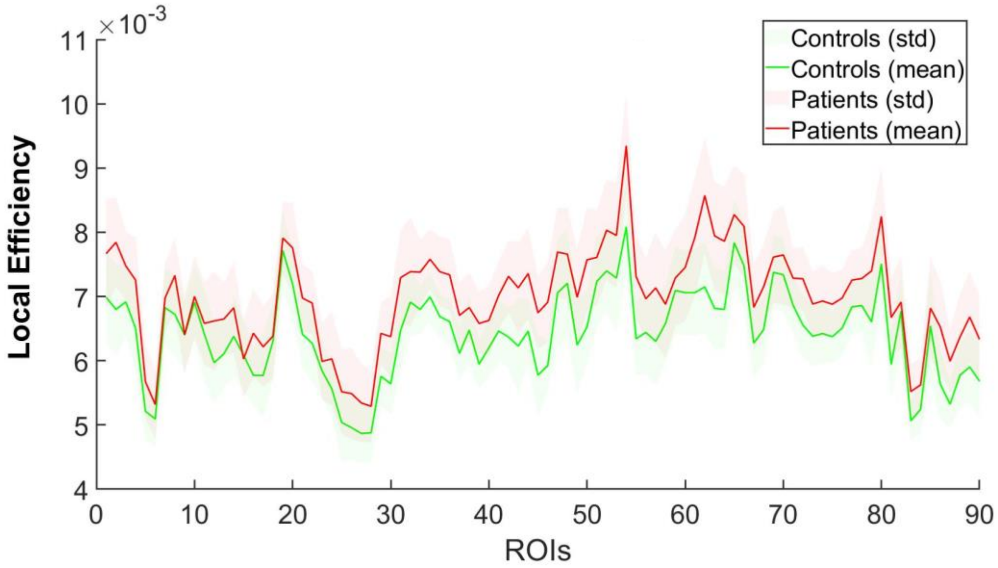


**Supplementary Fig. S2**: Local Efficiency at the *single-subject* level. Mean and standard deviation (std) of the values of local efficiency for each ROI in controls (green) and patients (red). The ROIs are listed following the order of the AAL atlas.

We found an overall increase in the PSD and in the network metrics comparing patients and controls, but the differences resulted significant only in the subset of the twelve ROIs reported in the paper. We also checked if there was a correlation between the results in the power spectra and the network metrics, no significant correlation was found between the power in delta and beta band and the network metrics. A correlation was found between the local efficiency and power in theta-alpha band, but it was not generalized to all the ROIs (Supplementary Fig. S3).

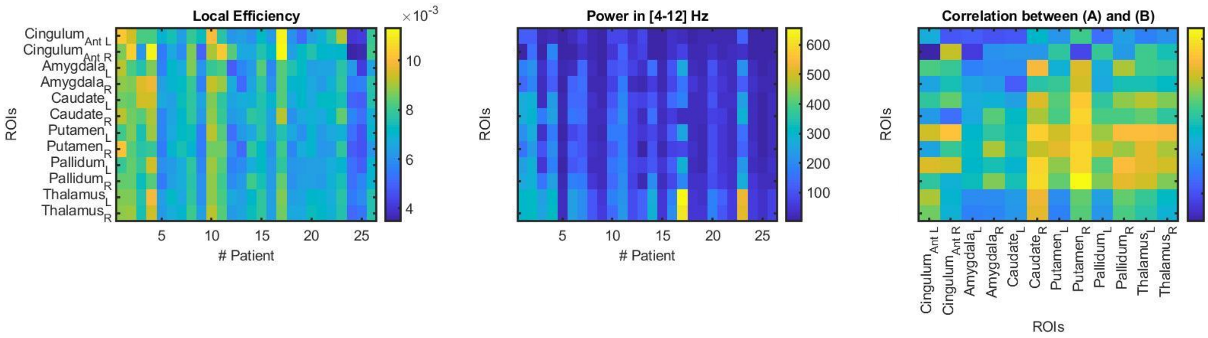


(a) (b) (c)

**Supplementary Fig. S3**: Correlation between the local efficiency and power in the twelve selected ROIs. (a) Values of local efficiency for each patient. The color code stands for the intensity of local efficiency; (b) Values of power in [4-12] Hz for each patient. The color code stands for the value of the power in $(\mu A/mm^3)^2$; (c) Correlation between the values of (a) local efficiency and the values of (b) power in [4-12] Hz in all the patients for each ROI. The color code stands for the r-value of correlation. All y -axes report the ROIs in the left (L) and right (R) hemispheres as labeled in (a), x-axes report the patient number or ROIs.
